# Protein Crowding Mediates Membrane Remodeling in Upstream ESCRT-induced Formation of Intraluminal Vesicles

**DOI:** 10.1101/834457

**Authors:** Susanne Liese, Eva Maria Wenzel, Ingrid Kjos, Rossana V. Rojas Molina, Sebastian W. Schultz, Andreas Brech, Harald Stenmark, Camilla Raiborg, Andreas Carlson

**Affiliations:** Department of Mathematics, Mechanics Division, University of Oslo, N-0851, Oslo, Norway; Department of Molecular Cell Biology, Institute for Cancer Research, Oslo University Hospital, Montebello, N-0379, Oslo, Norway

## Abstract

As part of the lysosomal degradation pathway, the endosomal sorting complexes required for transport (ESCRT-0-III/VPS4) sequester receptors at the endosome and simultaneously deform the membrane to generate intraluminal vesicles (ILVs). Whereas ESCRT-III/VPS4 have an established function in ILV formation, the role of upstream ESCRTs (0-II) in membrane shape remodeling is not understood. Combining experimental measurements and electron microscopy analysis of ESCRT-III depleted cells with a mathematical model, we show that upstream ESCRT-induced alteration of the Gaussian bending rigidity and their crowding on the membrane induces membrane deformation and facilitates ILV formation: upstream ESCRT-driven budding does not require ATP consumption as only a small energy barrier needs to be overcome. Our model predicts that ESCRTs do not become part of the ILV, but localize with a high density at the membrane neck, where the steep decline in the Gaussian curvature likely triggers ESCRT-III/VPS4 assembly to enable neck constriction and scission.

**Significance Statement:** Intraluminal vesicle (ILV) formation plays a crucial role in the attenuation of growth factor receptor signaling, which is mediated by the endosomal sorting complex required for transport (ESCRT-0-III/VPS4). The general dogma has been that the upstream ESCRTs (0-II) sequester the receptors at the surface of endosomes and the downstream ESCRTs (III/VPS4) remodel the endosome membrane leading to the abscission and formation of receptor-containing ILVs. We now show that the upstream ESCRTs not only sequester cargo, but in addition play an essential role for the initiation of membrane shape remodeling in ILV budding. Through a combination of mathematical modeling and experimental measurements we show that upstream ESCRTs facilitate ILV budding by crowding with a high density in the membrane neck region.

## Introduction

Sorting and compartmentalisation of biomaterials lie at the heart of cellular processes and play a fundamental role in the lysosomal degradation pathway to regulate cellular activities. Transmembrane proteins in the plasma membrane, such as growth factor receptors, are internalized by endocytosis and degraded in lysosomes [1]. As a part of this pathway, intraluminal vesicles (ILVs) with a typical diameter in the order of 50nm are formed inside endosomes [2] (Supplementary Figure S1). The ILV formation starts with a small deformation of the endosome membrane, which grows over time and finally leads to a cargo-containing vesicle within the endosome lumen. ILV generation is an essential part of the endocytic downregulation of activated receptors to ensure signal attenuation, failure of which can result in tumourigenesis [2, 3, 4, 5]. In addition, ILVs can reach the extracellular environment as exosomes, where they become signaling entities enabling intercellular communication. Altered exosomes can serve as tumor biomarkers [6, 7, 8]. In spite of the fundamental role ILVs play in the endocytic pathway, surprisingly little is known about the mechanochemical crosstalk that regulates their formation.

The biophysical process that leads to the formation of an ILV is in many ways inverse to clathrin-mediated endocytosis at the plasma membrane, as the membrane bud protrudes away from the cytosol (Fig. 1a) and the vesicle shape is not dictated by a protein scaffold [3]. Cargo sorting and ILV formation is mediated by the endosomal sorting complex required for transport (ESCRT) [4, 9, 10, 11, 12, 13], which consists of four sub-complexes, ESCRT-0, -I, -II and - III, and the accessory VPS4 complex. Each of the sub-complexes have different, yet complementary functions [4, 10]. ESCRT-0 contains binding domains for the endosome membrane, ubiquinated cargo and clathrin, which enables ESCRT-0 to sequester cargo material into patches, so-called microdomains, on the endosome membrane [14]. Interestingly, the role clathrin plays in ILV formation differs significantly from its role in endocytosis, as it promotes ILV formation [15], but does not form a basket-like scaffold. Instead, a rather flat clathrin coat is bound to the ESCRT microdomain [16, 17, 14, 18]. ESCRT-0 recruits ESCRT-I that leads to the recruitment of the complete ESCRT machinery. While ESCRT-0, -I, -II sequester transmembrane cargo proteins and facilitate ILV formation, ESCRT-III and the ATPase VPS4 enable constriction of the membrane neck leading to the formation of an ILV [19]. Notably, the only energy consuming step in the membrane remodeling process is the membrane scission, involving the ATPase VPS4 [20, 21]. This is especially remarkable, since the energy required for a flat lipid bilayer to form a spherical vesicle in absence of a protein coat, is orders of magnitude larger than the thermal energy, thus creating an energy barrier inhibiting vesicle formation [22, 23].

**Figure 1:**
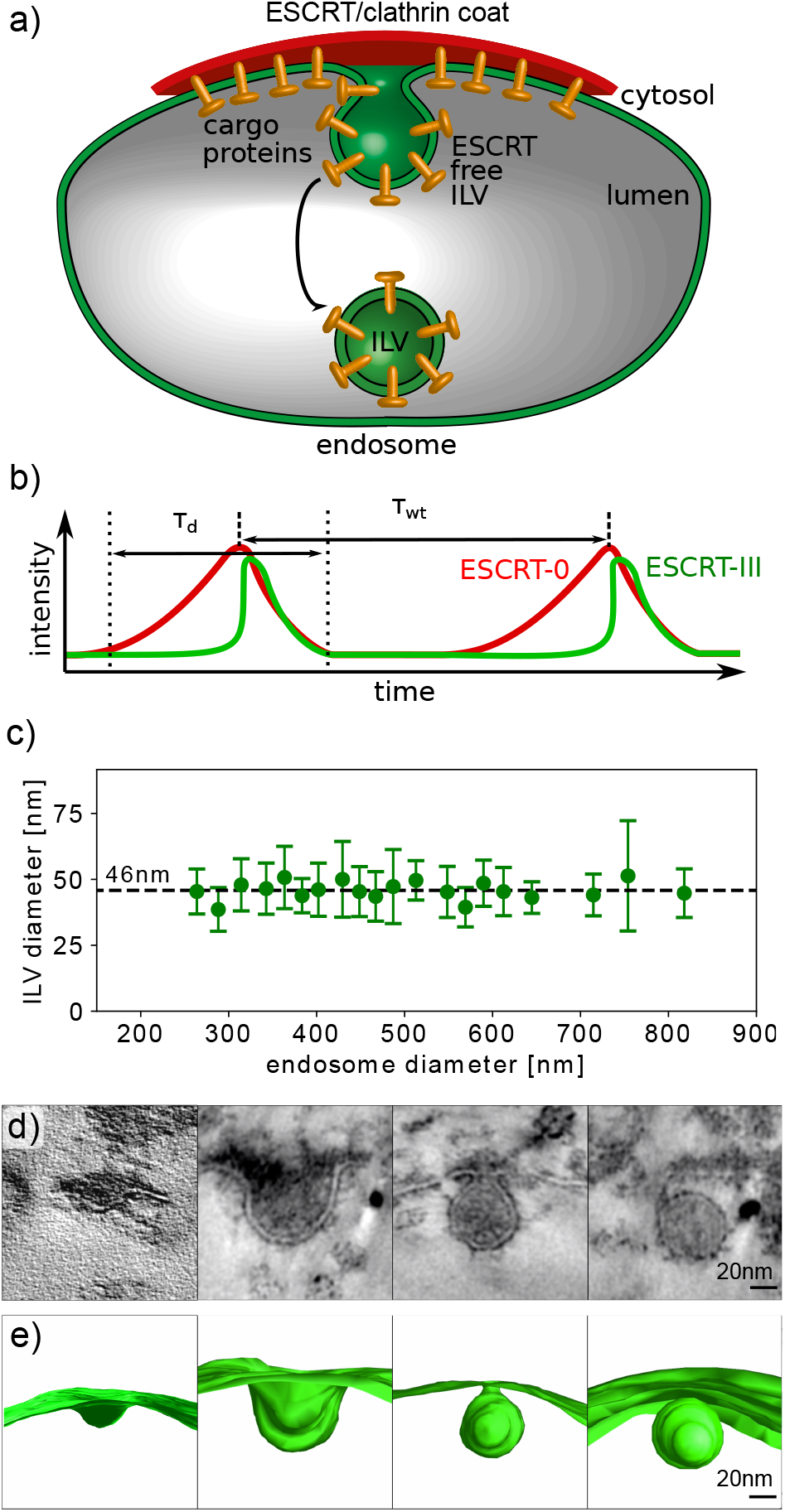
Illustration of the formation of ILVs: a) Schematic illustration of the formation of an ESCRT-free ILV. Transmembrane cargo proteins, ESCRTs and clathrin form a microdomain at the endosome, where cargo proteins are sorted by the ESCRT machinery into ILVs. b) We illustrate the characteristic time scales from the measured fluorescence intensity of ESCRT-0 and ESCRT-III [15]. The fluorescence signal intensity increases and decreases during the dwell time *τ*_d_. The mean waiting time between two consecutive ILV formation events is denoted by *τ*_wt_. c) Experimental measurements of 477 ILV diameters in HeLa cell endosomes exhibit a narrow distribution of ILV size independent of the endosome diameter. The ILV diameter was measured in TEM images. The average and the standard deviation are calculated over at least 10 ILV diameters. The endosome diameter is sorted into bins with a width of 20nm, the center of the bins is chosen such that the number of data points in each bin is maximized. d) During ILV formation the endosome membrane transitions through intermediate membrane shapes, illustrated by the TEM micrographs and classified (from left to right) into pit-shape, U-shape, Ω-shape and abscised vesicle [15]. e) We reconstruct the three dimensional membrane shapes, from TEM tomograms [15] (Supplementary Video 1). Subfigure d) and e) modified from [15]

Experiments have shown that the recruitment of ESCRTs to the endosome occur in a time periodic fashion [15, 24], where each recruitment cycle is associated with the formation of a single ILV. Transmission electron microscopy (TEM) images and electron tomography have also provided a high-resolution information of the membrane shapes during the budding process [15]. Today, *in*-*vivo* measurement techniques are not able to record the time evolution of fluorescent markers and the membrane shape simultaneously. To gain a detailed understanding of ESCRT assembly on the endosome membrane giant unilaminar vesicles (GUV) have been used as an *in*-*vitro* model system to study the formation of micrometer sized vesicles at their membrane [9]. The *in*-*vitro* experiments demonstrate that ESCRTs are enriched in the vesicle neck, but do not coat the main portion of the vesicle [9].

Among the ESCRT sub-complexes, ESCRT-III and VPS4 have received most attention in both experimental and theoretical studies, due to their fundamental role in a broad range of cellular processes involving membrane scission, e.g., cytokinesis and virus budding [25, 19, 26, 27, 28, 29, 30, 31, 32]. Theoretical and experimental studies have shown that the polymerization of ESCRT-III filaments into spirals promotes membrane buckling [33] and neck scaffolding [34], which in combination with the tension exerted by ESCRT-III, is suggested to cause closure of the membrane neck [35]. Much less is known about how the upstream ESCRTs affect membrane-remodeling, even though they play an as critical role in ILV formation [15, 36]. The lack of a description of the biophysical mechanisms generating membrane deformations at the endosome contrasts the large number of studies on clathrin mediated endocytosis at the plasma membrane, with similar membrane shapes, where theoretical modeling has provided invaluable information about the link between surface forces from transmembrane proteins [37], stochastic effects [38] and the resistive elastic forces [39, 40, 41, 42]. To observe ILV formation Rozycki et al. [43] assumed a uniform ESCRT coat on the membrane, which exhibits preferred binding to negative Gaussian curvature. Mercker et al. considered the influence of a spontaneous Gaussian curvature caused by the structure of the ESCRT-I and ESCRT-II supercomplex [44]. These models highlight that Gaussian bending is essential to understand the formation of an ILV.

Given our limited understanding of the biophysical mechanisms by which the upstream ESCRTs mediate ILV formation, we set out to investigate their role in this membrane remodeling process. By combining experimental measurements with theoretical modeling we answer the following questions:

1. What is the biomechanical mechanism that allows ILVs to form and does this process require an active force or a source of external energy?
2. How do ESCRT proteins organize at the endosome to later form an ESCRT-free vesicle?
3. How do the dynamics of ESCRT recruitment couple to the endosome membrane shape?

We find that ILV formation is facilitated by the upstream ESCRT proteins’ ability to alter the Gaussian bending rigidity and their crowding on the membrane. Finally, we verify our theoretical model by analyzing endosomal budding profiles in cells depleted of downstream ESCRTs.

## Results

### Underpinning experimental measurements

Fluorescence measurements of ESCRT protein dynamics were performed by live-cell microscopy of human cancer cells (HeLa), which show that ILV formation is accompanied by an oscillatory increase and decrease of the ESCRT concentration on the limiting membrane of endosomes [15], as illustrated in Fig.1b. In each recruitment cycle, the fluorescent signal of ESCRT-0 continuously increases over a time span of about three minutes, before it abruptly starts to decrease for about two minutes. In addition, it was shown that ESCRT-I and clathrin have similar dynamics as ESCRT-0. We therefore only need to consider the temporal evolution of ESCRT-0. The dynamic features of ESCRT-III are distinctly different from the ESCRT subunits 0-I. Once the ESCRT-0 signal reaches a maximum in fluorescence intensity, ESCRT-III exhibits a jump in its fluorescence intensity over just a few seconds, before it decreases with a decay time similar to ESCRT-0. The dynamics of ESCRT assembly are described by two time scales: the dwell time of ESCRT-0 *τ*_d_=(161±94)s and the periodicity of the ESCRT-recruitment cycle, or equivalently the mean waiting time *τ*_wt_ =(203±47)s [15].

TEM imaging of HeLa cancer cells reveals that ILVs have an average diameter of ≈46nm and that there is no apparent trend with respect to the endosome size, see Fig. 1c. A description of the membrane remodeling process that leads to ILV formation must therefore include a mechanism that is robust in terms of setting the vesicle diameter. TEM imaging also allows us to describe the membrane shape at different stages, where we categorize the membrane profiles into three specific shapes: pit-, U- and Ω-shape (Fig.1 d, e). As there is no overweight of samples in either of these categories [15], despite that these images are taken for different cells and at random time points within the ILV formation cycle, it suggests that the membrane deformation takes place in a continuous rather than a jump-like fashion over time.

### Energy barrier

Next, we use the fluorescent signal data from the experiments to extract information about the energy barrier that has to be overcome as an ILV forms. The magnitude of the energy barrier is crucial to classify whether ILV budding happens passively, *i.e.*, initiated by thermal fluctuations, or as an active process that requires energy consumption. To determine the energy barrier that is associated with the formation of a single ILV, we use a theory derived by Kim et al. [45] that relates the height of an energy barrier in a diffusive process to the ratio of two characteristic time scales *τ*_d_/*τ*_wt_. We deploy this theory for an energy landscape with the shape of a harmonic potential. In other words, we assume that the system has to overcome a single energy barrier with a magnitude Δ*E*_B_ to form an ILV. In a diffusive process the time to reach and cross this energy barrier once, *i.e.* the dwell time *τ*_d_ in the experimental measurements, scales with Δ*E*_B_ as *τ*_d_ ~ (4/3 − 16/45*β*Δ*E*_B_)*β*Δ*E*_B_, while the mean time that passes before the energy barrier is crossed a second time, *i.e.*, the waiting time *τ*_wt_ scales with Δ*E*_B_ as 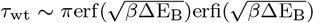, with *β*^−1^ the thermal energy. The ratio of the two experimental time scales, *τ*_d_/*τ*_wt_, provides an upper limit for the magnitude of the energy barrier Δ*E*_B_ ⪅0.6k_B_T (see Supplementary Figure S2). Since Δ*E*_B_ is of such low magnitude, it can be overcome by thermal fluctuations, giving us a first suggestion that ILV budding generated by the upstream ESCRTs is a passive process as we will further demonstrate below.

### Mathematical model

A mathematical model that describes ILV formation needs to incorporate how ESCRT proteins influence the shape of the endosome membrane. We consider ESCRT-0, -I, -II and clathrin as one effective complex that coats the endosome membrane (Fig. 2), and the membrane together with the embedded cargo proteins is treated as a homogenous elastic surface. Since the ILV size is much smaller than both the endosome (see Fig. 1c and Supplementary Figure S1) and the ESCRT microdomain (several hundred nanometers in diameter) [15], we simplify the model by considering only a part of the endosome membrane that is approximately planar (Fig. 2), where cargo proteins are evenly distributed within the domain. As part of the budding process a region within the ESCRT microdomain forms that is not coated by ESCRTs [9], which we define as the ESCRT-free region (Fig. 2). TEM tomography experiments show that ILVs are very close to being rotationally symmetric (Supplementary Video 1 and 2), which we adopt in the mathematical model, where the membrane is parameterized by the arc length *S* and the azimuthal angle *ψ* (Fig. 2). The *Z* coordinate and the radial coordinate *R* of the membrane contour are related to *S* and *ψ* through 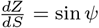, 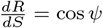. ILV formation involves a change in both mean and Gaussian curvature (see Supplementary Figure S3). A common feature among proteins that alter the Gaussian bending rigidity and promote membrane remodeling are *α*-helix motives in their secondary structure [46, 47]. The structural similarity between these proteins and the ESCRTs [48, 49, 50, 51] prompts us to model the Gaussian bending rigidity as dependent on the concentration of ESCRT proteins. The simplest mathematical description of a protein-induced Gaussian bending rigidity is to assume a linear response with respect to the ESCRT density *ρ*. The Gaussian bending energy Δ*E*_g_ then reads

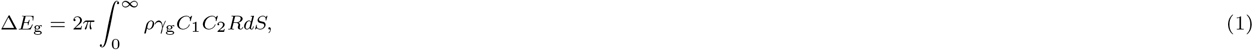

with the proportionality factor *γ*_g_ > 0. The two principle curvatures of the membrane are denoted as *C*_1_ = sin *ψ/R* and *C*_2_ = *dψ/dS*. In qualitative terms, a non-homogenous Gaussian bending rigidity describes the tendency of the membrane to deform into a neck-like shape, where a homogenous protein distribution (*ρ*=const.) would lead to a vanishing energy contribution, since the azimuthal angle is Ψ(*S* = 0) = 0 and Ψ(*S* → *∞*) = 0 at the inner and outer boundary and hence Δ*E*_g_ = 0 according to the Gauss-Bonnet theorem [52].

**Figure 2:**
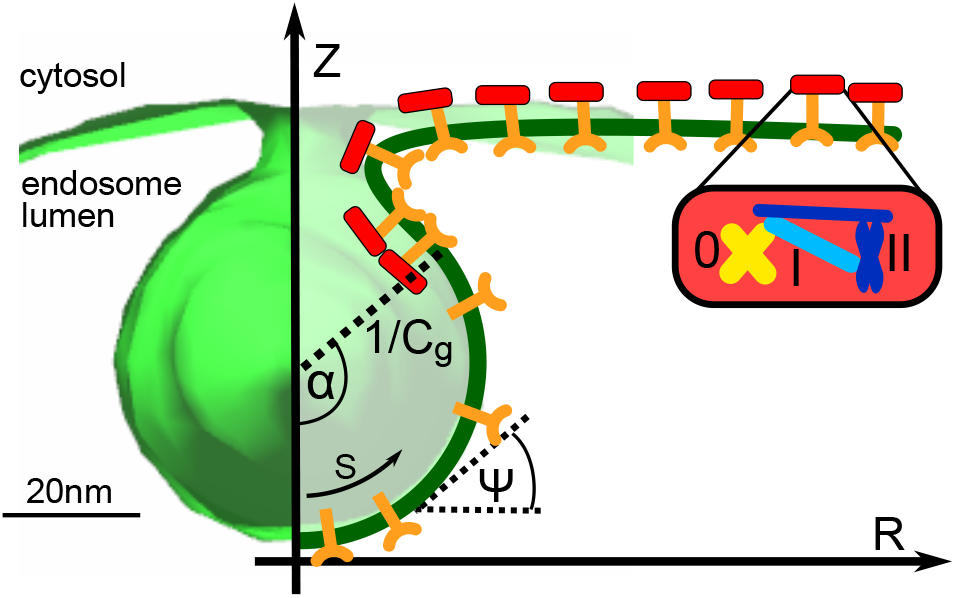
ILV parameterization: A part of the endosome membrane reconstructed from TEM micrographs (in green) is shown together with the parametrization of the membrane in the mathematical model. Membrane bound coat proteins (red blocks), which include ESCRT-0, -I, -II and clathrin, bind to the transmembrane proteins (Y-shaped orange markers). The endosome membrane deforms into an ESCRT-free spherical vesicle with a curvature *C*_g_, surrounded by a neck region with an elevated ESCRT density. The extent of the coat-free area is quantified by the opening angle *α*. The membrane shape is defined by the arc length *S* and the azimuthal angle *ψ*, where we treat the membrane as being axially symmetric around the Z-axis.

In addition to Δ*E*_g_, the total membrane energy has to account for membrane bending and stretching as well as protein crowding. By following the Helfrich model for lipid bilayers [22], we describe the membrane together with the embedded cargo proteins as a thin elastic sheet. The bending energy Δ*E*_*κ*_ is then obtained as an integral of the squared mean curvature over the entire surface

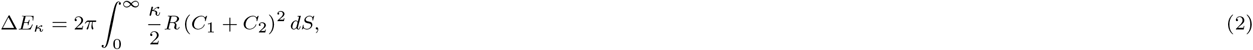

with the bending rigidity *κ*.

In the limit of an endosome that is much larger than the ILV, as is the case in this system, the pressure that acts across the lipid bilayer causes an effective far field tension, with *σ* the surface tension coefficient [53]. This means that remodeling the membrane away from a flat shape requires a surface energy Δ*E*_*σ*_

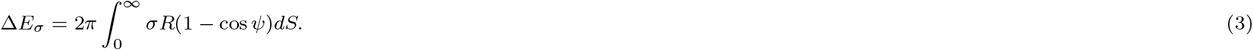

ESCRT proteins form a dynamic coat, *i.e.*, individual ESCRT proteins are continuously recruited to and dissociate from the endosome membrane [20, 54]. The binding energy Δ*E*_*μ*_ between the membrane and the coat is proportional to the local ESCRT density *ρ* and the coat area

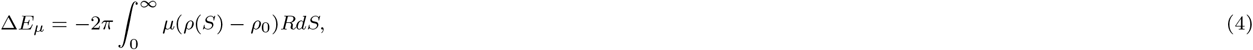

with *μ* the binding energy per unit area. The second term in Eq. 4 subtracts the binding energy of a flat continuously coated membrane with a uniform ESCRT density *ρ*_0_.

The amount of ESCRT proteins that can bind to the endosome membrane is limited, as they experience an effective steric repulsion [55, 56], primarily generated by volume exclusion, which we approximate up to second order in *ρ* by the second virial coefficient *ν*_2_. The corresponding steric repulsion energy Δ*E*_s_, is written as

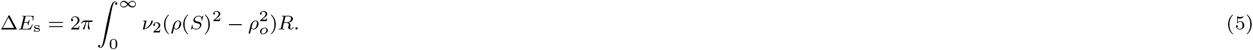

The second term in Eq. 5 subtracts the energy of a flat, uniformly coated membrane. Protein crowding has been shown theoretically and experimentally to be a mechanism that facilitates membrane deformation [56, 57]. Together, the binding energy (Eq. 4) and the steric repulsion (Eq. 5) represent the crowding effect that stem from the supercomplex of cargo proteins, ESCRTs and clathrin.

The total change in energy Δ*E* as we start from a flat membrane and progress towards a budding membrane, is at each stage in the process given by the sum of the five energy contributions

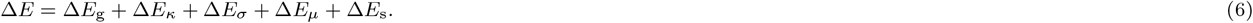

A non-uniform distribution of ESCRT proteins will lead to an additional energy term that penalizes large gradients in the density profile, which leads to an effective line tension at the boundary between the ESCRT-free and the coated region. Similarly, also protein diffusion along the membrane will smear the protein density profile. However, this line tension contribution is a very small correction to the membrane energy (Supplementary Figure S6) and is neglected here. A spontaneous curvature induced by a protein-scaffold is not expected to play a significant role in ILV budding as *in – vitro* experiments using GUVs have shown that ESCRTs do not coat ILVs [9]. In addition, a spontaneous curvature induced by cargo proteins can be disregarded as a cause for membrane shape remodeling, since ILVs serve as a sorting compartment for a large variety of transmembrane proteins and there is no biophysical indication that these proteins all induce a similar spontaneous curvature [58].

The ratio of the specific binding energy *μ* and the proportionality factor of the Gaussian bending rigidity *γ*_g_ defines an inverse length scale *C*_g_ that we use to express the membrane energy in dimensionless variables,

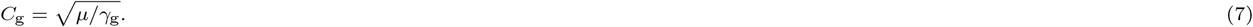

We will show below that *C*_g_ corresponds to the mean curvature of the ESCRT-free ILV. The ratio of *μ* and the second virial coefficient *ν*_2_ defines a density *ρ*_0_, which is the baseline ESCRT density on the flat membrane 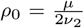. Furthermore, we introduce the non-dimensional numbers 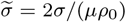 and *ϵ* = *ρ*_0_*γ*_g_/*κ* = *μγ*_g_/(2*ν*_2_*κ*), which dictate the energy landscape of the membrane remodeling process. 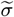 describes the ratio between surface tension and binding energy, which increases for a high surface tension and for a small *μ* or a low *ρ*_0_, *i.e.*, for a weak interaction between the ESCRT proteins and the endosome membrane or for a low ESCRT density on the non-deformed membrane. The tension *σ* in biological membranes can vary quite significantly *σ* ∩ 10^−6^ – 10^−3^N/m [59, 60, 61]. The interaction energy between biological membranes and proteins is typically in the range of *μ* ≈1k_B_T [62]. The size of ESCRT proteins is in the order of 10nm [50, 63, 51], which in turn leads to an estimate for the ESCRT density of *ρ*_0_ ≈0.01 nm^−2^. Combining these data gives us the expected physiological range of the dimensionless number 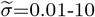. *ϵ* describes the ratio of bending rigidities associated with the mean and the Gaussian curvature, which are typically in the same order of magnitude [64] and we expect 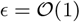.

### ESCRT density on the endosome

The ESCRT proteins form a stable microdomain at the endosome for several minutes. However, individual ESCRT proteins are rapidly exchanged within a few seconds [54, 20], which suggests that the ESCRTs can quickly adapt their local density to changes in the membrane curvature in order to minimize the overall energy. By minimizing Eq. 6 with respect to the ESCRT density *ρ* gives us a relation between *ρ* and the principal curvatures

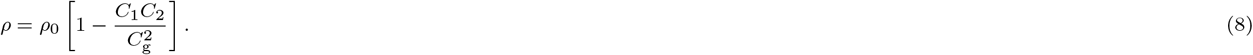

We see from Eq. 8 that the ESCRT density is uniform with *ρ* = *ρ*_0_ when *C*_1_ = 0 or *C*_2_ = 0, *i.e.*, on a flat membrane. In contrast, the protein density is reduced where the membrane exhibits a positive curvature, *i.e.*, at the center of the budding vesicle, while *ρ* increases in regions of negative Gaussian curvature, *i.e.*, in the neck region of the vesicle. Eq. 8 reveals that an ESCRT-free membrane bud, with *ρ* = 0, follows from a spherical membrane shape, where both principal curvatures are given by *C*_1_ = *C*_2_ = *C*_g_. The curvature *C*_g_ of the ESCRT-free region (Eq. 7) is determined by a balance between binding energy and the Gaussian bending rigidity. To gain a deeper understanding of the parameters that determine the vesicle curvature, we consider the individual contributions to the membrane energy Eq. 2-6. There are two terms that can generate a negative contribution to the energy: the binding energy Δ*E*_*μ*_ and the Gaussian bending energy Δ*E*_g_. While Δ*E*_g_ = 0 for a uniform coat of upstream ESCRTs, the system gains energy in the form of a Gaussian bending energy, if an ESCRT-free membrane bud forms, since Δ*E*_g_ = 0 in the coat-free region and Δ*E*_g_ < 0 in the outer region, where the membrane exhibits a negative Gaussian curvature. At the same time, the system pays an energetic penalty due to a reduced binding energy. It is the balance between binding energy and Gaussian bending rigidity that determines the curvature of the ESCRT-free ILV.

### Endosome membrane shapes and ESCRT density profiles

The membrane shape and energy depend only on the curvature *C*_g_ and the dimensionless numbers *ϵ* and 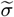. In Fig. 1c and [15] we determine an average ILV diameter of 46nm, which is equivalent to a curvature *C*_g_ ≈0.04 nm^−1^. Defining the curvature *C*_g_ from experimental measurements enables a reduction of the number of free parameters in our model to just two, *ϵ* and 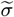.

If the membrane adopts a spherical shape with a mean curvature *C*_g_, the ESCRT-free area is given by 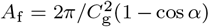, with *α* the angle formed between the tip of the bud at the *Z*-axes and the start of the ESCRT coated region as illustrated in Fig. 2. To understand how the membrane energy changes as this region increases, we minimize the total energy Eq. 6 under the constraint that *α* is fixed. We have fixed *ϵ*=2.0 and 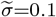, which leads to an energy barrier that is similar to our analytical estimate based on the characteristic ESCRT-recruitment time scales, Δ*E*_B_ ≈0.6k_B_T. Fig. 3 shows the quasi-static membrane shape predicted by our mathematical model and the ESCRT density together with the experimental membrane shapes. From the comparison between the mathematical model and the experimental data we determine the opening angle *α*=0.35*π*, 0.5*π*, 0.7*π*, corresponding to pit-shape, U-shape and Ω-shape of the membrane.

**Figure 3:**
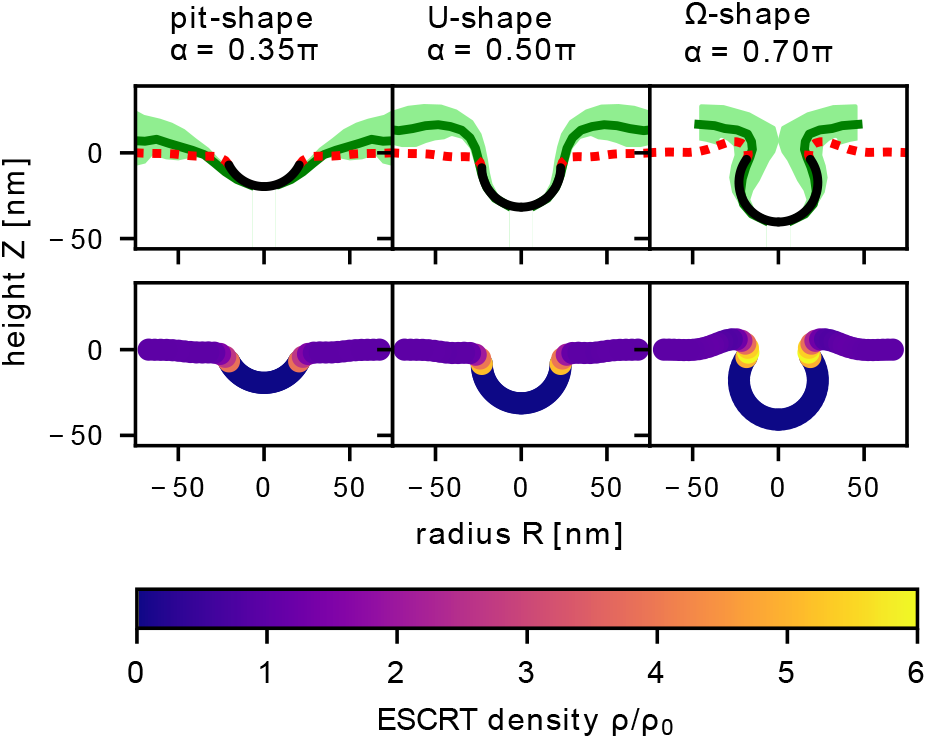
Vesicle budding: We compare the experimentally measured endosome shapes with the minimal energy shapes (Eq. 6) for different angles *α*, when *ϵ* = 2 and 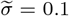. (top row) The experimental membrane shapes [15] are grouped into three categories: pit-shape (*α* = 0.35*π*), U-shape (*α* = 0.5*π*) and Ω-shape (*α* = 0.7*π*). For each subgroup the average shape is shown as a solid green line, while the standard deviation is indicated by the shaded area. The ESCRT-free vesicle bud is shown by the solid black line, while the coated region is shown by the red dashed line. (bottom row) The ESCRT density is shown along the membrane for the three characteristic shapes, which exhibits an elevated ESCRT density in the neck region.

We turn next to see if the model can also help to understand the *in – vitro* experimental observations based on GUVs, which have also demonstrated ESCRT-mediated membrane budding in the absence of ESCRT-III. The size of the vesicles formed in these experiments, with a typical vesicle diameter in the *μ*m range, differs largely from ILVs [9]. According to Eq. 7 the size difference stems from a variation of the specific binding energy *μ* or the Gaussian bending rigidity, quantified by the proportionality factor *γ*_g_. We expect the capability of ESCRT proteins to induce a Gaussian bending rigidity to be similar in both the *in-vivo* and *in-vitro* system, giving nearly the same *γ*_g_. The variation in vesicle size is therefore attributed to different values of the specific binding energy *μ*, which we suspect is lower in GUV experiments as they lack cargo proteins. A significant increase of the ESCRT density was found in the vesicle neck region on GUVs [9], which is also predicted by our theoretical model (Fig. 3). We observe that when the membrane has an Ω-shape, the ESCRT density profile has a pronounced maximum in the neck region that is six times greater than *ρ*_0_.

### Upstream ESCRT-mediated membrane deformation does not require energy input

In Fig. 4a) we show how the membrane energy depends on *α* as we go from a flat membrane (*α* = 0) to an Ω-shaped membrane (*α/π* → 1) for three different combinations of *ϵ* and 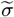, where *ϵ*=2.0 and 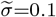 correspond to physiologically realistic values, while *ϵ*=1.3, 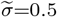 and*ϵ*=0.6 and 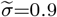 result in an energy barrier that cannot be overcome by thermal fluctuations and thus prevents ILV formation. For *ϵ*=2.0, 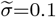 and a vesicle diameter of 46nm, the opening angle of the energy barrier (*α* = 0.04*π*) corresponds to an ESCRT-free area of just 26nm^2^. Hence, only a few ESCRT-proteins have to desorb to overcome the energy barrier and initiate the budding of an ILV.To better understand which part of the energy dominates during the shape transition we show the individual contributions to the energy together with the total membrane energy for *ϵ*=2.0, 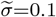 in Fig. 4b). The change in both binding energy (Eq. 4) and surface tension (Eq. 3) are small compared to the overall change in energy. The energy generated by the steric repulsion of proteins (Eq. 5) and the bending of the membrane (Eq. 2) both increase continuously, thus opposing ILV formation. The only sizeable negative contribution comes from the Gaussian bending term (Eq. 1), which dominates the total membrane energy as the membrane shape approaches scission (*α/π* → 1). The divergence of the Gaussian energy as the angle approaches *α/π* → 1 is a characteristic of the large local variation of the Gaussian bending rigidity. Theoretical studies have shown that a partially formed vesicle and a finite vesicle neck is not stable, *i.e.*, the energy diverges, if the variation in Gaussian bending rigidity is large compared to the bending rigidity associated with mean curvature [65].

**Figure 4:**
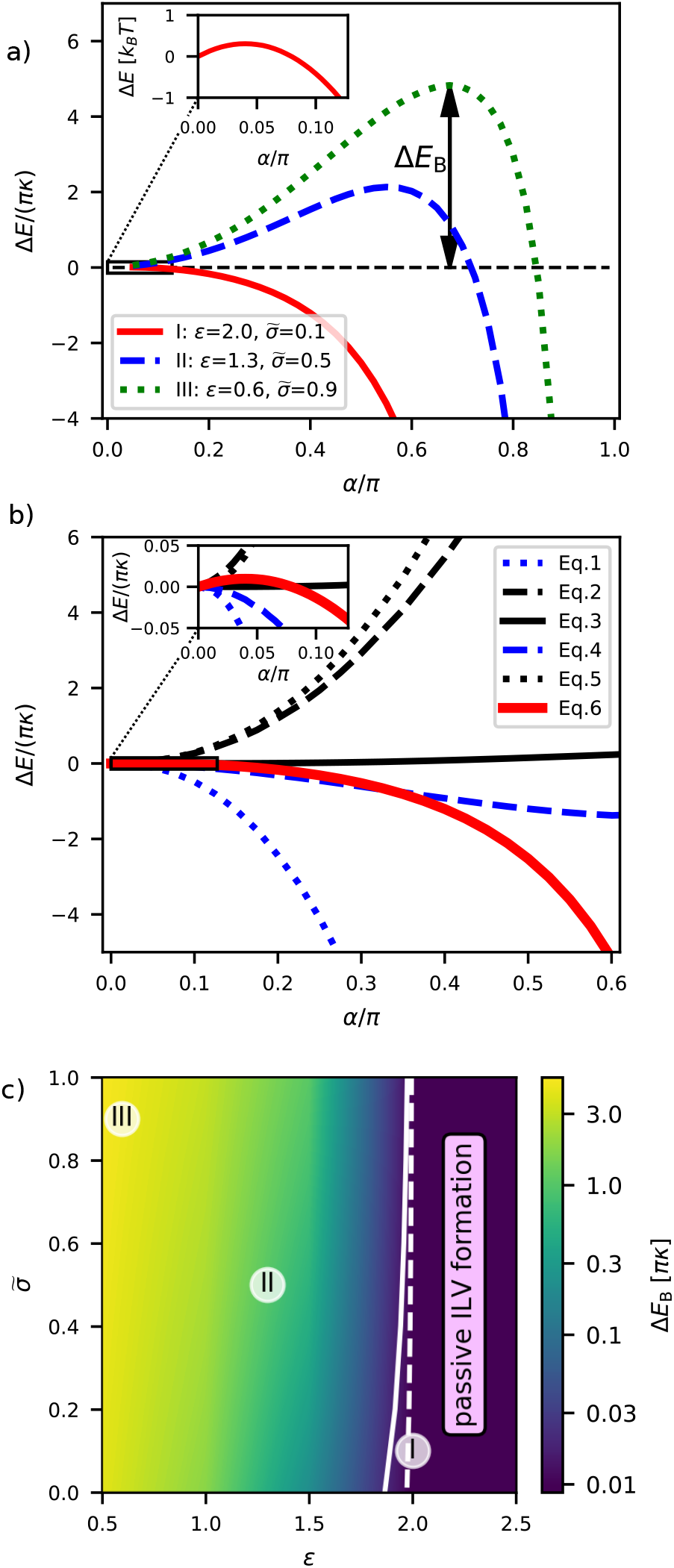
Energy barrier: a) To understand how the energy changes with the shape changes of the membrane, we plot Δ*E*_B_ as a function of the opening angle *α* for (I) *ϵ* =2.0, 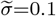; (II) *ϵ* =1.3, 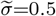 and (III) *ϵ* =0.6, 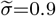. The energy landscape (I), solid red line, corresponds to the membrane shape transition shown in Fig. 3. Inset: The maximum of the energy for *ϵ* =2.0, 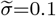 is found around *α/π*=0.04. To convert the energy into units of k_B_T, the bending rigidity is set to *κ*=10k_B_T [23]. b) We plot the different contributions of Δ*E* (Eq. 1-5) together with the total energy Eq. 6 for (I) *ϵ*=2.0, 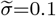 to help illustrate which part produces a positive (inhibiting ILV formation) and negative (promoting ILV formation) energy. It is clear that both steric repulsion of proteins and bending of the membrane is energetically costly, while the Gaussian bending energy becomes negative and grows in magnitude as ILV formation takes place (*α/π* → 1). c) We scan the phase space in *ϵ* (ratio of the bending rigidities associated with the mean and the Gaussian curvature) and 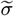 (ratio of surface tension and binding energy) in numerical simulations to determine their influence on the energy barrier Δ*E*_B_. The magnitude of the energy barrier is shown as a color map with a logarithmic scale. The threshold energy barrier for passive ILV formation is illustrated by the solid line when Δ*E*_B_=2.0k_B_T and the analytical estimate for the energy barrier of ILV formation, Δ*E*_B_=0.6k_B_T is shown by a dashed line.

Next, we scan the phase space of *ϵ* ∈ [0.5–2.5] and 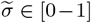 in numerical simulations, where we minimize the energy (Eq. 6) to determine the magnitude of the energy barrier Δ*E*_B_ as a function of *α*. In Fig. 4c), we see that Δ*E*_B_ decreases with increasing *ϵ* and decreasing 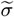. In Fig. 4c) we illustrate the region where an ILV is formed passively, by drawing a solid line for Δ*E*_B_=2k_B_T, which for a bending rigidity of *κ*=10k_B_T [23] is equivalent to Δ*E*_B_/(*πκ*) ≈0.06 in dimensionless units. In addition, we show a dashed line for Δ*E*_B_=0.6k_B_T as determined by our analyzes as an estimate for the energy barrier of ILV formation. The phase space in Fig. 4c) shows that passive ILV formation is feasible for a wide range of values for 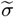 and *ϵ*. Our mathematical model is thus robust and implies that upstream ESCRT-mediated membrane deformation does not require energy.

### Neck closure

*In – vitro* experiments on GUVs have shown that ESCRT-III and Vsp4 alone are sufficient to induce vesicle formation with an inverse topology, similar to *in – vivo* ILV budding [66]. However, in ILV formation both upstream ESCRTs (ESCRT-0, -I, -II) and ESCRT-III as well as Vsp4 and cargo proteins are present, which play different and crucial roles in cargo sorting and ILV formation. The mathematical description of the ILV formation puts us in the position to relate the ESCRT recruitment dynamics, which we have tracked experimentally using ESCRT-0 fluorescence, with the transient membrane shapes. We determine the amount of excess ESCRT proteins (Δ*n*) as the integral over the protein density on the membrane, where the ESCRT density exceeds the baseline value *ρ*_0_. In rescaled units Δ*n* reads as

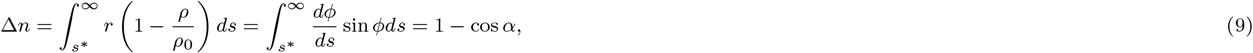

with *s*\* the arc length where the ESCRT coated region begins. Inserting Eq. 8 into the left hand side of Eq. 9, we find that Δ*n* is expressed as an integral over *ϕ*, where we know the angle at the inner and outer boundary, with *ϕ*(*s*\*) = *α* and *ϕ*(*s* → ∞) = 0. While the membrane undergoes a shape transition from a flat surface to an Ω-shape, Δ*n* appears to increase continuously. The experimental measurement of the fluorescent signal of ESCRT-0 is proportional to Δ*n*. Hence, Eq. 9 enables us to determine the shape evolution in terms of the time dependency of *α* from the fluorescent ESCRT-0 signal (Methods section). The corresponding time evolution of *α*, shown in Fig. 5b, reveals a continuous rather than a jump-like vesicle formation in accordance with the experimental observations (Fig. 5a and [15]).

**Figure 5:**
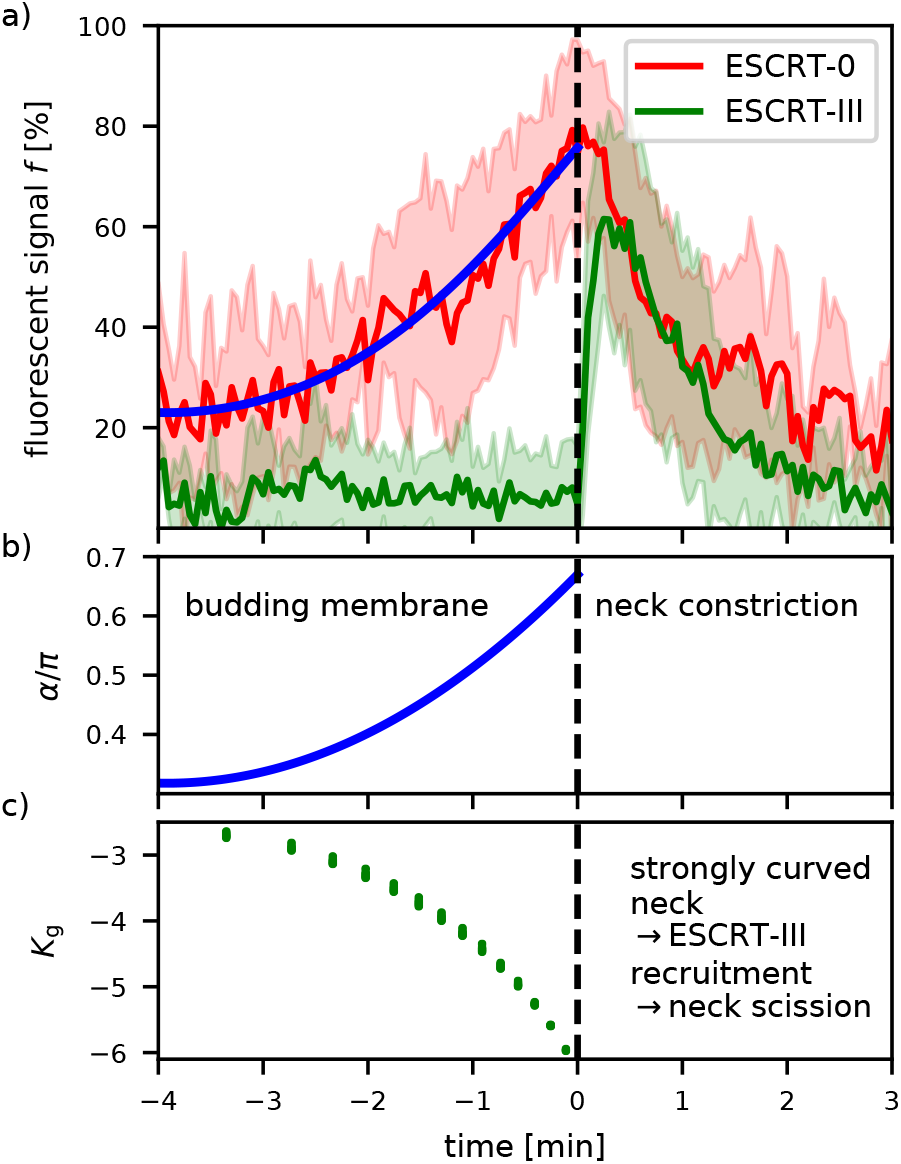
ESCRT-III recruitment: a) The experimental signal intensity of fluorescently labeled HRS (ESCRT-0) and CHMP4B (ESCRT-III) is shown as a function of time. The average over 23 isolated signal curves is shown together with the standard deviation (shaded area). The fluorescent signal of ESCRT-0 is proportional to the amount of adsorbed proteins, which allows us correlate the magnitude of the fluorescent signal with the opening angle *α* (Methods Section). Figure modified from [15] b) By combining the experimentally measured time evolution of the fluorescent marker with the theoretically predicted membrane shapes, we determine the time evolution of *α* according to the fit in subfigure a. c) The Gaussian curvature in the neck, *i.e.*, at the boundary between ESCRT-free and - coated region, is determined for six different combinations of *ϵ*, 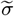, with *ϵ* ∈ [2.0, 2.5] and 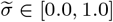. The Gaussian curvature strongly increases in magnitude as the membrane transitions from a flat to an Ω-shape.

ESCRT-III mediated scission of the membrane neck is the final step of ILV formation and ESCRT-III exhibits a preferred binding to curved regions of the membrane [27]. To quantify the membrane curvature in the ILV neck, we define the magnitude of the rescaled Gaussian curvature as 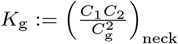 at the boundary between ESCRT-free and - coated region. Fig. 5c shows *K*_g_ for six combinations of 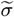 and *ϵ*, with *ϵ* ∈ [2.0, 2.5] and 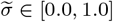, *i.e*, in the regime of passive ILV formation, where we see that all data points collapse on to a single curve exhibiting the same behavior. It illustrates that even though the energy barrier strongly depends on *ϵ*, the Gaussian curvature is a key feature of the membrane shape depending only marginally on the parameters 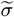 and *ϵ*, *i.e.*, the membrane shape is robust towards fluctuations in membrane tension and bending rigidity. By comparing the fluorescent signal of ESCRT-III and the Gaussian curvature in the neck region, we speculate that the steep decrease of the Gaussian curvature *K*_g_, triggers the assembly of ESCRT-III and subsequently leads to membrane scission forming an ILV.

### Experimental verification of the theoretical predictions

Our mathematical model implies that the increase in upstream ESCRT-density on the endosome can drive membrane deformation in the absence of energy input, and define the size and the shape of the forming ILV. Moreover, our model implies that the steep decrease in the Gaussian curvature obtained by the accumulation of the upstream ESCRTs triggers the observed abrupt recruitment of ESCRT-III/VPS4, leading to neck constriction and scission. Our model thus predicts to observe inward membrane buds in the endosomal membrane in the absence of ESCRT-III. To verify the theoretical predictions experimentally, we depleted ESCRT-III components, which also precludes the recruitment of the VPS4 ATPase [67], testing our hypothesis that upstream ESCRTs i.e. ESCRT-0, -I and -II, suffice to deform the limiting membrane of endosomes. siRNA-mediated knockdown of the ESCRT-III components CHMP2A and CHMP4 led to a pronounced depletion of these proteins as verified by Western blotting (Supplemantary Figure S7a). Immunofluorescence experiments showed that ESCRT-III depletion led to enlarged late endosomes and a hyperrecruitment of HRS to endosomes, as expected when the ESCRT machinery is perturbed (Supplementary Figure S7b) [68, 69, 70]. We interpret these findings as indicative of a good knockdown efficiency. Importantly, the epidermal growth factor (EGF) could still be endocytosed and reach endosomal compartments in ESCRT-III depleted cells (Supplementary Figure ??b), allowing us to study the ILV process in depth, in newly formed EGF-induced endosomes. To investigate whether the limiting membrane of the endosome can be deformed in the absence of ESCRT-III, we performed electron microscopy on ESCRT-III depleted cells. To mark newly formed endosomes, we prebound an antibody recognizing the extracellular part of epidermal growth factor receptor (EGFR) and added a secondary antibody conjugated to 10 nm gold particles, keeping the cells during the whole labelling procedure at 4°C to stall endocytosis. Then we stimulated endocytic uptake of EGFR by incubating cells with EGF ligand for 12 minutes at 37°C before high-pressure freezing, freeze substitution and electron microscopy. At 12 minutes of EGF stimulation, ESCRTs were shown previously to be very active in ILV formation [15]. From these electron microscopy sections, we counted the number of formed ILVs relative to the endosome area. ESCRT-III depletion resulted in a reduced number of abscised ILVs when compared to non-targeting control, see (Supplementary Figure S7c,d,e), indicating that the overall ILV formation was impaired. Importantly, we were still able to observe forming ILV buds in ESCRT-III depleted cells (siControl: 21, siCHMP4: 19, siCHMP2A: 19 from 47, 47 and 31 endosomes, respectively), supporting the notion that buds do form in the absence of ESCRT-III. Given the lower number in abscised ILVs, but the unchanged or even slightly increased number in buds (Fig. 6), we speculated that ESCRT-III depletion might stall or slow down the progression of ILV formation.

**Figure 6:**
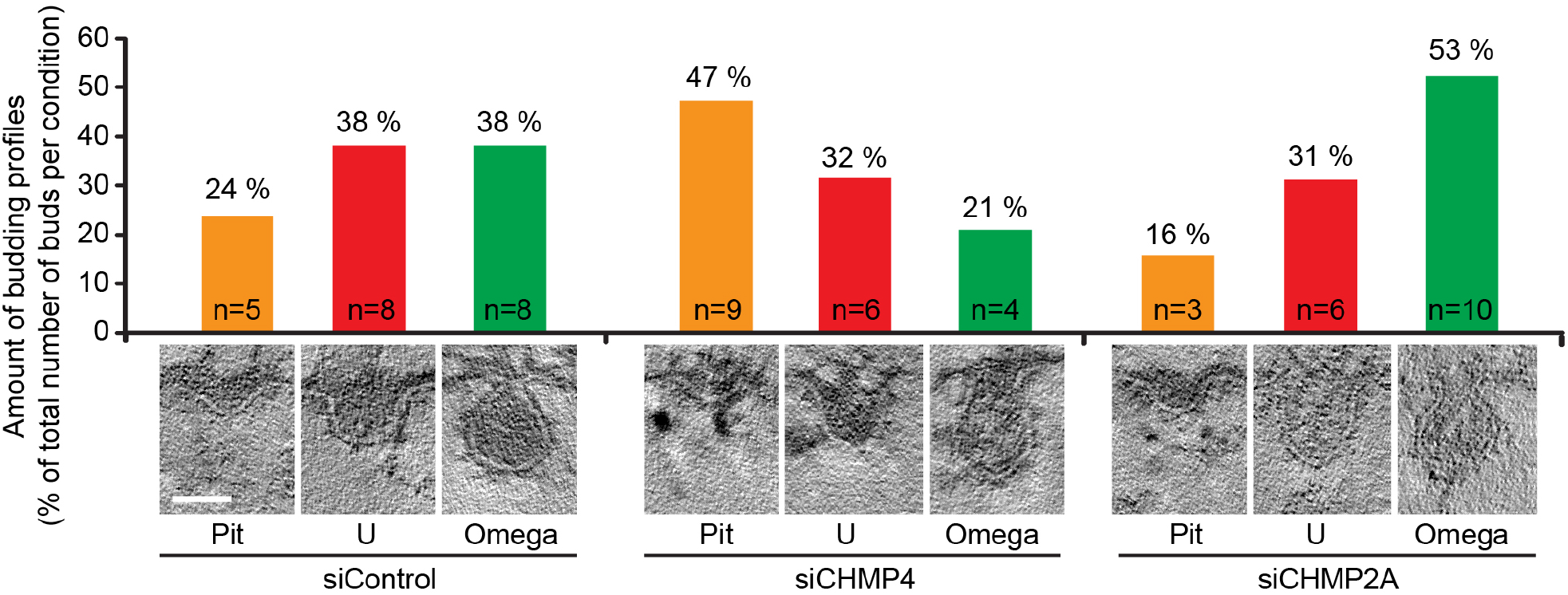
Formation of membrane buds in control and ESCRT-III depleted cells: HeLa cells were transfected with control RNA or siRNA targeting CHMP4 isoforms or CHMP2A for 48 hours, stimulated with EGF for 12 min to internalize gold labeled EGFR, and analyzed by electron microscopy as described in the methods. EGFR-gold positive endosomes were collected from tomograms of individual 200 nm thick EM sections and all detectable budding profiles in the limiting membrane of the endosomes were identified and organized into pit-, U- and ‘Ω’-shapes by measuring the ratio between the neck with and the bud depth (with/depth <0.7 ‘pit’; 0.7-1.7 ‘U’; >1.7 ‘Ω’). The graph represents the amount of profiles in the different categories in %, n indicates the number of profiles observed in total. The micrographs show representative membrane shapes. Scale bar 40 nm.

To analyse the stages of ILV formation, we sorted the budding profiles according to their morphology and observed a reduction in the number of constricted buds (Ω-shapes) in the absence of CHMP4 (Fig. 6), which is considered to be the dominating subunit in ESCRT-III filaments [71]. Importantly, in the absence of CHMP4 filaments, the limiting membrane of endosomes showed an accumulation of non-constricted invaginations, likely mediated by the upstream ESCRT machinery (ESCRT-0, -I and -II). Depletion of CHMP2A, which is the most downstream ESCRT-III filament and which serves to recruit the ATPase VPS4 [67], resulted in a higher proportion of omega-shaped buds when compared to control or CHMP4 knockdown, arguing for a stalled process of ILV formation at a late stage directly before scission. To summarize, our experimental data supports the findings from our mathematical model, where the upstream ESCRT machinery starts the membrane deformation process through protein crowding, likely involving transmembrane cargo proteins and this does not require energy in the form of ATP.

## Discussion

By combining mathematical modeling and cell biological data we are able to point to the biophysical determinants that facilitate ILV budding by upstream ESCRTs, which is a consequence of the interplay between ESCRT-induced Gaussian bending rigidity and their crowding on the membrane. Our mathematical model highlights that while ESCRT dissociation at the budding site is biologically desirable, since it enables the cell to reuse the ESCRT proteins, it is also a physical prerequisite to form an ILV as the systems benefits from the negative energy from Gaussian bending only if the Gaussian bending rigidity, or equivalently the ESCRT-density, is non-homogenously distributed at the membrane. The ILV size is set by a balance between the loss of binding energy in the ESCRT-free region and by the increase of the Gaussian bending energy in the neck region. Our model predicts; (i) a high density of ESCRTs in the membrane neck, (ii) the shape of the endosome membrane during budding (pit-, U- and Ω-shape), (iii) vesicle formation to be a continuous process in time, all in accordance with experimental observations.

By treating the experimental time scales for recruitment of ESCRTs as a diffusive process we analytically predict the energy barrier the system must overcome to form a vesicle Δ*E*_B_ ≈ 0.6k_B_T. Furthermore, our mathematical model that accounts for membrane bending, binding and crowding of proteins, and the spatial distribution of the upstream ESCRTs, shows that membrane budding is a self-organized passive process, that does not need ATP consumption, which explains, why it is sufficient for the ATPase VPS4 to bind to the ESCRT complex in the final stage of membrane constriction and scission [19]. By scanning the phase space in the ratio of bending rigidities (*ϵ*) and the ratio of the surface tension and the binding energy 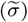 in numerical simulations, we show that ILVs may form passively over a wide range of parameters. Thus, to inhibit membrane budding the system must be perturbed such that the energy barrier exceeds beyond the range that can be affected by thermal fluctuations. We predict that a change in membrane tension by a hypertonic shock (changing 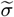), would suppress ILV formation. The energy landscape is also sensitive to changes of the bending rigidities (changing *ϵ*), which leads us to speculate about the role of the clathrin layer that is bound to the ESCRT microdomain. The physical properties of the clathrin layer are not yet understood. We have shown earlier that ILV formation is severely impaired in the absence of clathrin recruitment to endosomes [15], which we can now rationalize in the framework of our model, where absence of clathrin binding corresponds to an effective decrease of the binding energy *μ* (*i.e. ϵ* is reduced) and stalls ILV formation through a higher energy barrier.

We used the predicted ESCRT density from the quasi-static theoretical model to correlate it to the increase in the experimentally measured fluorescent intensity of ESCRT-0, which gives a prediction of the opening angle *α* over time. Our analysis point to a continuous rather than jump-like transition from a pit-shape, to a U-shape and an Ω-shape, in concordance with the fact that there is no overweight in samples for either of these shape categories found in the TEM images of different endosomes and at random time points in the budding process. Together this leads us to argue that upstream ESCRTs play a crucial role not only in sequestering cargo proteins, but also in the membrane shape remodeling in accordance with earlier work on GUVs [9]. Importantly, our data points to the upstream ESCRTs as determinants for the initial membrane budding: Fluorescence microscopy data shows that ESCRT-0 and ESCRT-I get enriched at the endosome membrane over several minutes. On a flat membrane this would lead to a significant increase in steric repulsion between the ESCRT proteins and an energy increase. Instead, the system evades an energetic penalty by forming a membrane bud in the pathway to ILV formation. Moreover, our electron microscopy data shows that budding profiles indeed form in the absence of ESCRT-III, underscoring the role of upstream ESCRTs in the budding process.

The membrane shape transition is accompanied by a steep decline in the Gaussian curvature in the neck region, which we believe is a trigger for ESCRT-III assembly to facilitate membrane scission, since ESCRT-III binds preferentially to negatively curved membranes [27]. Qualitatively similar ESCRT-III recruitment dynamics have been found in HIV budding [72], where Gag proteins assemble on the membrane over a longer time span, while ESCRT-III shows a spike-like recruitment over time. HIV budding, where the Gag proteins cause an effective spontaneous curvature [73, 72], resembles ESCRT-induced ILV budding morphologically. In particular, the formation of a curved membrane neck is likely a prerequisite in both cases to promote ESCRT-III assembly [27]. Upon assembly, the different subunits of the ESCRT-III complex are expected to have differential functions in the membrane shaping process. Whereas CHMP4B is the main component of the ESCRT-III filament that oligomerizes in the membrane neck and recruits the complete ESCRT-III machinery, CHMP2A functions to recruit the ATPase VPS4 [12, 67, 71]. This notion is supported by our experimental data, where the shapes of the budding profiles that can be observed in CHMP4 versus CHMP2A depleted cells are different: CHMP2A depletion caused an accumulation of constricted membrane buds as expected, consistent with a failure to mediate membrane scission in the absence of VPS4. In CHMP4 depleted cells, however, we observed an accumulation of un-constricted budding profiles, mainly pit-shapes. This finding is interesting, as it suggests that CHMP4 not only functions in membrane constriction and in the recruitment of the complete ESCRT-III machinery and thus VPS4. In addition, CHMP4 filaments could have a stabilizing role which aids in the transition to a constricted form. In the absence of CHMP4 filaments, membrane buds will continuously form due to the crowding of the upstream ESCRTs, but in the absence of a stabilizing factor, some profiles might revert to a pit-shape.

Together our observations show that the experimentally measured increase in the fluorescent signal of upstream ESCRTs is a hallmark of the change in membrane shape at the endosome. This implicates the upstream ESCRTs together with clathrin and cargo proteins in the membrane remodeling process and adding to their role as cargo sorting molecules. The generic nature of protein crowding and a spatially varying Gaussian bending rigidity on cell membranes suggests that the developed model can have implications beyond understanding budding of ILVs by upstream ESCRTs, as it may also help understand other membrane remodeling processes.

## Material and Methods

### Cell culture and siRNA transfections

HeLa (Kyoto) cells (obtained from D. Gerlich, Insitute of Molecular Biotechnology, Wien, Austria) were grown according to ATCC guidelines in DMEM high Glucose (Sigma-Aldrich) supplemented with 10% fetal calf serum, 100 U ml-1 penicillin, 100 *μ*g ml-1 streptomycin and maintained at 37°C under 5% CO_2_. The cell line is authenticated by genotyping and regularly tested for mycoplasma contamination. Cells were transfected using Lipofectamine RNAiMax transfection reagent (Life Technologies) following the manufacturer’s instructions. All siRNAs were purchased from Ambion^®^ (Thermo Fisher Scientific) and contained the Silencer Select modification. The following siRNA sequences were used: CHMP4B#1 (5’-CAUCGAGUUCCAGCGGGAGtt-3’) CHMP4B#2 (5’-AGAAAGAAGAGGAGGACGtt-3’), CHMP4A (5’-CCCUGGAGUUUCAGCGUGAtt-3’), CHMP4C (5’-AAUCGAAUCCAGAGAGAAAtt-3’), CHMP2A (5’-AAGAUGAAGAGGAGAGUGAtt-3’). Experiments were performed 48 h after transfection. Non-targeting control Silencer Select siRNA (predesigned, catalogue number 4390844) was used as a control. The first EM experiment was done with 50 nM siCHMP4B#2, in the other two EM experiments a cotransfection of CHMP4B#2, siCHMP4A, siCHMP4C (25 nM each) was done to avoid potential compensation of CHMP4A and CHMP4C. We did not notice a more penetrant phenotype in the triple knockdown compared to depletion of CHMP4B only. 50 nM siCHMP2A was used for both experiments.

### Antibodies and reagents

Antibodies: rabbit-anti-CHMP2A (10477-1-AP from Proteintech, immunofluorescence 1:500), mouse anti–*β*-actin (A5316 from Sigma-Aldrich, Western blot 1:10000), rabbit-anti-CHMP4A (sc-67229 from Santa Cruz, Western blot 1:500), rabbit anti-CHMP4B was generated as described previously [74] (Western blot 1:1000), rabbit anti-HRS (immunofluorescence 1:100) has been described previously [17], mouse anti-LAMP1 (H4A3 from Developmental Studies Hybridoma Bank, immunofluorescence 1:1000), mouse anti-EGFR (555996 from Pharmingen, extracellular labeling of EGFR). All secondary antibodies used for immunofluorescence studies were obtained from Jacksons ImmunoResearch Laboratories or from Molecular Probes (Life Technologies). Secondary antibodies used for western blotting where obtained from LI-COR Biosciences GmbH.

### Immunoblotting

Cells were washed with ice-cold PBS and lysed with NP40 lysis buffer (0.1% NP40, 50 mM Tris-HCl pH7.5, 100 mM NaCl, 0.2 mM EDTA, 10% Glycerol) supplemented with “Complete EDTA-free protease inhibitor cocktail” (05056489001 from Roche). Cell lysates were spun for 5 min. at 16000 g at 4°C to remove nuclei. The supernatant was mixed with 4x SDS sample buffer (200 mM Tris-HCl pH 6.8, 8% SDS, 0.4% Bromphenol blue, 40% Glycerol) supplemented with DTT (0.1 M endconcentration) and then subjected to SDS-PAGE on 12% or 4–20% gradient gels (mini-PROTEAN TGX; Bio-Rad). Proteins were transferred to PVDF membranes (TransBlot^®^ TurboTM LF PVDF, Bio-Rad) followed by antibody incubation in 2% BSA in Tris-buffered saline with 0.1% Tween20. Membranes incubated with fluorescent secondary antibodies (IRDye680 or IRDye800; LI-COR) were developed with an Odyssey infrared scanner (LI-COR).

### Immunostaining and confocal fluorescence microscopy

Cells grown on coverslips were incubated with 50 ng ml-1 EGF-Al647 (E35351, Thermo Fisher Scientific) for 10 min at 37°C. Cells were then placed on ice and permeabilized with ice-cold PEM buffer (80 mM K-Pipes, pH 6.8, 5 mM EGTA, and 1 mM MgCl_2_) supplemented with 0.05% saponin (S7900-25g from Merck Life Science) for 5 min on ice to decrease the fluorescent signal from the cytosolic pool of proteins before fixation in 3% formaldehyde for 15 min [75]. Cells were washed twice in PBS and once in PBS containing 0.05% saponin before staining with the indicated primary antibodies for 1 h. After washing three times in 0.05% saponin in PBS, cells were stained with secondary antibodies for 1 h, and washed three times in PBS. The cells were mounted in Mowiol containing 2 mg ml-1 Hoechst 33342 (Sigma-Aldrich). Confocal fluorescence microscopy was done with a Zeiss LSM780 microscope (Carl Zeiss MicroImaging GmbH) using standard filter sets and laser lines and a Plan Apo 63x 1.4 N.A. oil lens. All images within one dataset were taken at fixed intensity settings below saturation.

### Electron microscopy and measurements

HeLa cells were grown on poly-l-lysine coated sapphire discs. To label newly internalized EGFR following EGF-stimulation, cells were first washed with ice cold PBS and incubated on ice with an antibody recognizing the extracellular part of EGFR (mouse anti-EGFR, Pharmingen). After washing four times with ice cold PBS, cells were incubated with Protein A-10 nm gold conjugate (UMC Utrecht Dept. of Cell Biology) which recognizes the Fc portion of the mouse IgG2b primary antibody. Cells were again washed four times with ice cold PBS and then stimulated with 50 ng ml-1 EGF in warm DMEM for 12 minutes before high pressure freezing was done. Sapphire discs were high pressure frozen using a Leica HPM100, and freeze substitution was performed according to the following schemes. a) sapphire discs were transferred to sample carriers containing freeze substitution medium (0.1% (w v-1) uranyl acetate in acetone, 1% H_2_O) and placed in a Leica AFS2 for automated freeze substitution according to the following program: −90°C for 48 h, temperature increase to −45°C for 9 h, −45 for 5 h, 3x wash with acetone, temperature increase from −45°C to −35°C, Lowicryl infiltration with stepwise increase in Lowicryl HM20 concentration (10%, 25%, 75%, 4 h each) and concomitant temperature increase to −25°C, 3x 10 h with 100% Lowicryl HM20, UV-polymerization for 48 h, temperature increase to +20°C, UV-polymerization for 24 h. b) sapphire discs were transferred to cryo-vials containing freeze substitution medium (0.5% uranyl acetate, 1% OsO4 and 0.25% glutaraldehyde in acetone) and placed in a Leica AFS2 for automated freeze substitution according to the following program: −90°C for 12 h, temperature increase to −60°C for 6 h, −60°C for 2 h, temperature increase to −20°C for 30 min, temperature increase from −20°C to 4°C for 15 min. The samples were left at 4°C for 30 min before transfer to room temperature, 3x wash in acetone and epon embedding with stepwise increase in epon concentration (25%, 50%, 75%, 100%). 200 nm sections were cut on an Ultracut UCT ultramicrotome (Leica, Germany) and collected on formvar coated slot grids. Samples were imaged using a Thermo ScientificTM TalosTM F200C microscope equipped with a Ceta 16M camera. Single-axes tomography tilt-series were recorded between −60° and 60° tilt angles with 2° increment. Tomograms were computed in IMOD using weighted back projection [76]. Measurements of endosome areas and ILV numbers were done in FIJI [77].

### Energy minimization

We rescale all lengths with *C*_g_ giving the dimensionless variables: *s* = *C*_g_ · *S*, *r* = *C*_g_ · *R*, *ψ*(*S*) → *ϕ*(*s*). By introducing this scaling of the variables we rewrite Eq. 6 in dimensionless form as

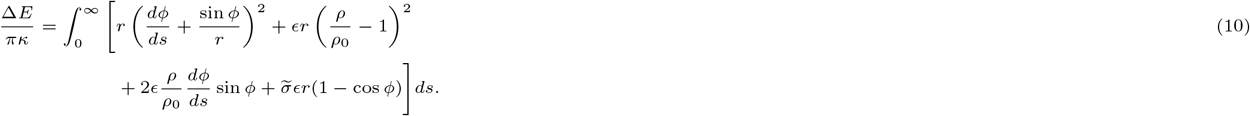

For any angle *α* the membrane shape in the inner region is described by a spherical cap with rescales mean curvature 1. The shape as well as the energy contribution in rescaled units are hence obtained analytically as *r* = sin *ϕ*, *z* = 1 − cos *ϕ* and 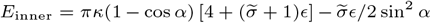. To minimize the total energy in the outer region numerically, we define a total arc length *s*_end_ at which the angle *ϕ* reaches zero. We set *s*_end_ = 15 to approximate the limit of an infinitely large surface. Similar to the method described by Rozycki et al. [43] we describe the angle *ϕ* in the outer region as a truncated Fourier series:

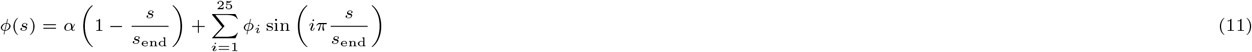

The radius *r* is then obtained from the relation *dr/ds* = cos *ϕ*. The prefactors *ϕ*_*i*_ are obtained by minimizing the energy Eq. 10 using the python *basinhopping* routine [78] (Supplementary information).

### Fluorescent signal fit

The intensity of the experimentally measured fluorescent signal *f* is assumed to be proportional to Δ*n*. To relate both quantities, we recall that the membrane shapes, which are closest to a fully formed ILV are Ω-shaped with an angle *α*=0.7*π*. We therefore assume the maximum of the fluorescent signal to correspond to Δ*n*\* = Δ*n*(*α* = 0.7*π*) = 1.59, which allows to predict how *α* varies in time in the experiment by fitting the fluorescent signal (Fig. 5a) to

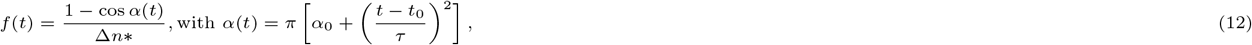

with the fit parameters *α*_0_=0.32, *t*_0_=−3.9min, *τ* =6.6min, where *τ* describes the characteristic time over which the fluorescent signal increases and *α*_0_ together with *t*_0_ account for an offset due to the background noise of the fluorescent signal.

## Acknowledgement

We thank Marianne Smestad and Ulrikke Dahl Brinch for expert help with electron microscopy sample processing and Else Munthe for technical assistance with the knock down experiments. S. L., R. R. M. and A. C. gratefully acknowledge funding from the Research Council of Norway Project Grant 263056.

