## Supplementary Information for "Protein Crowding Mediates Membrane Remodeling in Upstream ESCRT-induced Formation of Intraluminal Vesicles"

#### Endocytic Pathway

In Fig. S1 the role of ILVs within the endocytic pathway is shown schematically.

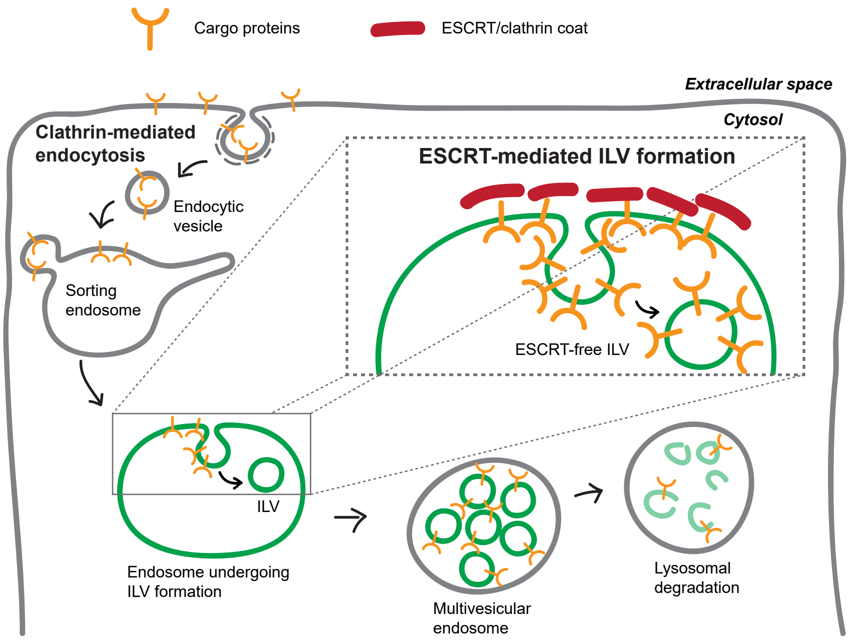


**Figure S1: Endocytic pathway** Transmembrane proteins (cargo), for example activated growth factor receptors, can enter the degradative endocytic pathway via endocytosis. To allow degradation in the lysosomes by lysosomal digestive enzymes, the cargo needs to be internalized into intraluminal vesicles (ILVs). Cargo sorting and the formation of ILVs is mediated by the ESCRT machinery, giving rise to multivesicular endosomes.

### Energy Barrier

For a diffusive process in one-dimension in a harmonic potential $E$ of the form $E(x) = 4\Delta E_\mathrm{B}\frac{x}{L}\left(1-\frac{x}{L}\right)$, with the energy barrier $E_\mathrm{B}$, the diffusion coordinate $x$ and the maximal value of the diffusion coordinate $L$, Kim et al. derived the following expressions for the transition path time $\tau_\mathrm{d}$, *i.e.* the time to cross the barrier if the process starts at $x$=0 and does not return to $x$=0, and the waiting time $\tau_\mathrm{wt}$, *i.e.* the mean time to cross the barrier, when starting from $x$=0, where the process may return to $x$=0 multiple times [1]:

$$\tau_\mathrm{d} = \frac{L^2}{4D}F_{2,2}\left(\frac{\Delta E_\mathrm{B}}{k_\mathrm{B}T}\right) - \frac{L^2}{2D\sqrt{\pi\frac{\Delta E_\mathrm{B}}{k_\mathrm{B}T}}\,\mathrm{erf}\left(\sqrt{\frac{\Delta \mathrm{E_B}}{\mathrm{k_B T}}}\right)}\int_0^{\frac{\Delta E_\mathrm{B}}{\mathrm{k_B}T}} y^2 e^{-y^2}F_{2,2}(-y^2)dy, \tag{S1}$$

$$\tau_\mathrm{wt} = \frac{L^2}{D}\frac{\pi\mathrm{erf}(\sqrt{\Delta \mathrm{E_B/k_B T}})\mathrm{erfi}(\sqrt{\Delta \mathrm{E_B/k_B T}})}{8\Delta E_\mathrm{B}/k_\mathrm{B}T} \tag{S2}$$

with $F_{2,2}(y) = F_{2,2}(\{1,1\};3/2,2;y)$ the generalized hypergeometric function and $D$ the diffusion constant. In the limit of small and large energy barriers Eq. S1 is approximated by the analytic expression:

$$\tau_\mathrm{d} \approx \frac{L^2}{D}\left[\frac{1}{6} - \frac{2}{45}\frac{\Delta E_\mathrm{B}}{k_\mathrm{B}T}\right], \qquad \frac{\Delta E_\mathrm{B}}{k_\mathrm{B}T} \ll 1 \tag{S3a}$$

$$\tau_\mathrm{d} \approx \frac{L^2}{D}\frac{\ln\left(2e^{\gamma_e}\frac{\Delta E_\mathrm{B}}{k_\mathrm{B}T}\right)}{8\frac{\Delta E_\mathrm{B}}{k_\mathrm{B}T}}, \text{with } \gamma_e \approx 0.577 \text{ the Euler gamma constant} \qquad \frac{\Delta E_\mathrm{B}}{k_\mathrm{B}T} \gg 1 \tag{S3b}$$

Eq. S1 and Eq. S2 show that the ratio $\tau_\mathrm{d}/\tau_\mathrm{wt}$ only depends on the energy barrier $\Delta E_\mathrm{B}$. Fig. S2 shows the ratio $\tau_\mathrm{d}/\tau_\mathrm{wt}$ as a function of the energy barrier. For a small energy barrier the curve approaches as value of $\tau_\mathrm{d}/\tau_\mathrm{wt}$=1/3. The experimentally obtained values for the dwell time and the waiting time are: $\tau_\mathrm{d}$=(161±94)s, $\tau_\mathrm{wt}$(203±47)s [2], which leads to a ratio 0.3≤ $\tau_\mathrm{d}/\tau_\mathrm{wt}$ ≤1.6 (indicated as gray shaded are in Fig. S2 ). Hence the upper limit exceeds the maximal theoretical value of 1/3, which we attribute to the shape of the energy landscape, which is not a perfectly harmonic potential for the case of ILV formation. Within the limit of the experimental uncertainty, we obtain an upper limit for the energy barrier of $\Delta E_\mathrm{B}$ ≈0.6$\mathrm{k_B T}$ from the point where the theoretical curve (black line in Fig. S2) leaves the grey area.

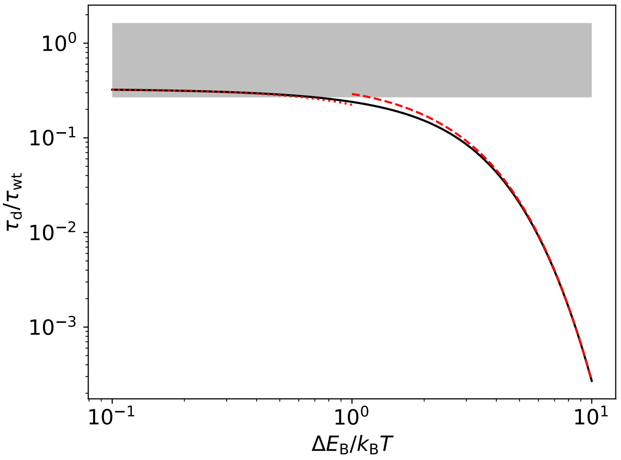


**Figure S2: Energy barrier** The ratio of transition path time and waiting time in dependence of the energy barrier $\Delta E_\mathrm{B}$ (Eq. S1, S2 solid black line, Eq. S3a, S2 dotted, red line and Eq. S3b, S2 dashed, red line). The range of experimental values is indicated as gray shaded area.

### Mean and Gaussian Curvature

A key assumption in our theoretical model is that ESCRTs induce a concentration dependent Gaussian bending rigidity. The membrane energy therefore depends on the mean curvature as well as on the Gaussian curvature. To gain a better understanding on how mean and

Gaussian curvature proceed along the membrane shape, as the membrane transitions from a flat surface to a spherical vesicle, Fig. S3 shows both curvatures for three exemplary membrane shapes. The same shapes are shown in Fig. 3 in the main text.

We rescale both mean and Gaussian curvature with the ILV curvature $C_\mathrm{g}$:

$$\text{Mean curvature: } \kappa_\mathrm{m} = \frac{\frac{d\psi}{dS} + \frac{\sin\psi}{R}}{2C_\mathrm{g}} \tag{S4}$$

$$\text{Gaussian curvature: } \kappa_\mathrm{g} = \frac{\frac{d\psi}{dS}\frac{\sin\psi}{R}}{C_\mathrm{g}^2} \tag{S5}$$

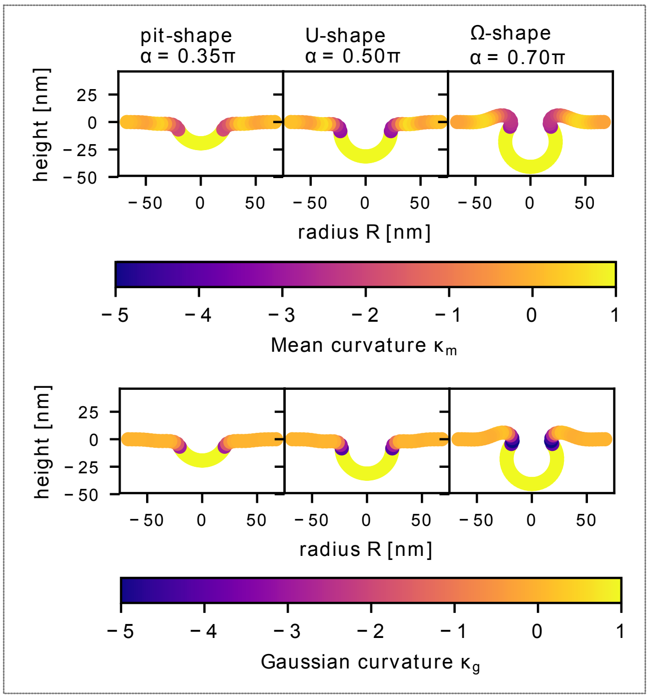


**Figure S3: Mean and Gaussian Curvature** The membrane shape is shown for different angles $\alpha$, corresponding to the three categories: pit-shape ($\alpha = 0.35\pi$), U-shape ($\alpha = 0.5\pi$) and Ω-shape ($\alpha = 0.7\pi$). The top row shows the mean curvature along the membrane shapes. The bottom row shows the Gaussian curvature along the membrane shapes.

### Numerical Method

The minimal energy shape of the outer region is obtained by parametrizing the angle $\phi$ as

$$\phi(s) = \alpha\left(1 - \frac{s}{s_\mathrm{end}}\right) + \sum_{i=1}^{N} \phi_i \sin\left(i\pi\frac{s}{s_\mathrm{end}}\right) \tag{S6}$$

The prefactors $\phi_i$ are obtained by minimizing the energy, Eq. 6 in the main text, using the python *basinhopping* routine [3] with 100 iteration steps and an initial step size of 0.01.

To optimize the number of terms $N$, we perform an energy minimization considering only the bending energy (Eq. 2 in the main text). Evidently the energy is minimal and equal to zero, if the mean curvature is zero. Hence, a membrane only subjected to bending energy is a suitable test case to compare the numerically and the analytically obtained minimal energy shape.

In Fig. S4 the bending energy of the outer membrane region is shown in dependence of the angle $\alpha$ for different $N$. As initial guess, we set the prefactors to $\phi_1$=-1.6 and $\phi_i$=0 for $i$ >1. With increasing $N$ the energy curve approaches the analytic result of $E_\kappa$=0. $N$=30 does not lead to a significant improvement compared to $N$=25. For numerical efficiency we hence chose $N$=25 for all numerical calculations.

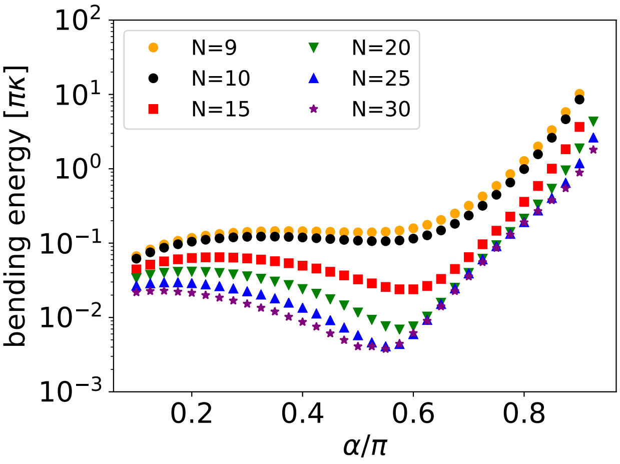


**Figure S4: Bending Energy** The numerically obtained bending energy of the outer membrane region. For a zero-mean-curvature shape the bending energy is zero, independent of the angle $\alpha$.

For all $N$ the numerically obtained bending energy deviates from $E_\kappa$=0 as $\alpha$ approaches $\pi$. Hence, the bending energy is over estimated around $\alpha = \pi$. However, the energy curves investigated in the main text show a maximum for much smaller $\alpha$, *i.e.* numerical error around $\alpha = \pi$ do not influence our findings reported in the main text.

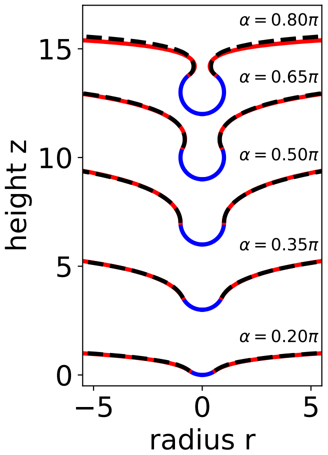


**Figure S5: Membrane Shape** Numerically obtained minimal energy shape (dashed, black line) and zero-mean-curvature shape (solid, red line). The shapes are shifted in $z$-direction for better visibility.

As a further test, we compare in Fig. S5 the numerically obtained minimal energy shape with a zero-mean-curvature shape for $N$=25 and different $\alpha$, which shows good agreement. To determine the zero-mean-curvature shape we solve the following initial value problem with the python *odeint* routine [3]:

$$\frac{d\phi}{ds} = -\frac{\sin\phi}{r} \qquad \phi(0) = \alpha \tag{S7}$$

$$\frac{dr}{ds} = \cos\phi \qquad r(0) = \sin\alpha \tag{S8}$$

$$\frac{dz}{ds} = \sin\phi \qquad z(0) = 1 - \cos\alpha \tag{S9}$$

The minimization of the full energy function (Eq. 6 in the main text) is performed in two steps. First, an energy minimization is performed considering only the bending energy. The thus obtained prefactors $\phi_i$ serve as an initial guess for the second step, where the full energy function is minimized.

### Line Tension

A non-uniform protein density leads to an effective surface tension, which acts to flatten the density profile:

$$\Delta E_\lambda = 2\pi \int_0^\infty dS \lambda \left(\frac{d\rho}{dS}\right)^2 \approx 2\pi\lambda\rho_0 2 \left(\frac{C_1(S^*)C_2(S^*)}{C_\mathrm{g}}\right)^2 R(S^*), \tag{S10}$$

with the proportionality factore $\lambda$. Since the largest density gradient is found at the boundary between the protein coated and uncoated region, the term $\left(\frac{d\rho}{dS}\right)^2$ is well approximated by a delta-distribution around $S^*$, with $S^*$ the arc length at the boundary of the coat-free vesicle. Hence, $\Delta E_\lambda$ acts like a line tension energy with an effective line tension $\lambda\rho_0^2$ that is scaled by the Gaussian curvature at the boundary between coated and uncoated region. Mercker et al. estimated the line tension of ESCRT proteins in the range of $\lambda\rho_0^2$ ≈0.005pN [4], which at about two orders of magnitude smaller than the typical line tension of lipid rafts [5]. Assuming this value for the line tension was motivated mainly by the observation that ESCRT-I and ESCRT-II do not form large clusters [6], which indicated a weak interaction among proteins.

To evaluate the impact of the line tension on the energy landscape, we write Eq. S10 in rescaled units as:

$$\frac{\Delta E_\lambda}{\pi\kappa} = \widetilde{\lambda} \left[r\,(c_1c_2)^2\right]_{s=s^*}, \text{ with } \widetilde{\lambda} = \frac{2\lambda\rho_0^2}{C_\mathrm{g}\kappa} \tag{S11}$$

and $c_1 = C_1/C_\mathrm{g}$, $c_2 = C_2/C_\mathrm{g}$. The rescaled radius at the boundary between coated and uncoated region is obtained analytically as $r(s^*) = \sin\alpha$, whereas the Gaussian curvature $c_1c_2$ is obtained from the energy minimization for $\widetilde{\lambda}$=0. This approximation serves as an upper limit of the energy term $\Delta E_\lambda$. In Fig, S6 the total energy is shown in dependence of $\alpha$ for $\epsilon$=2.0 and $\widetilde{\sigma}$=0.1. For a curvature of $C_\mathrm{g}$=(23nm)$^{-1}$, a bending rigidity $\kappa$=10k$_\mathrm{B}$T [7] and a line tension $\lambda\rho_0^2$=0.005pN the rescaled line tension becomes $\widetilde{\lambda}$=0.006. Fig. S6 shows that the energy landscape is not significantly altered for $\widetilde{\lambda}$=0.006.

If we assume an effective line tension that is several orders of magnitude larger, the line tension hinders the formation of a vesicle, which seems counter intuitive at first, since the radius at the boundary between coated and uncoated region decreases as the membrane transforms from a U-shape to an Ω-shape. However, the line tension is scaled by the squared of the Gaussian curvature at $s = s^*$, which increases rapidly with increasing $\alpha$ as a consequence, the term $\widetilde{\lambda}\left[(c_1c_2)^2\right]_{s=s^*}$ opposes neck closure.

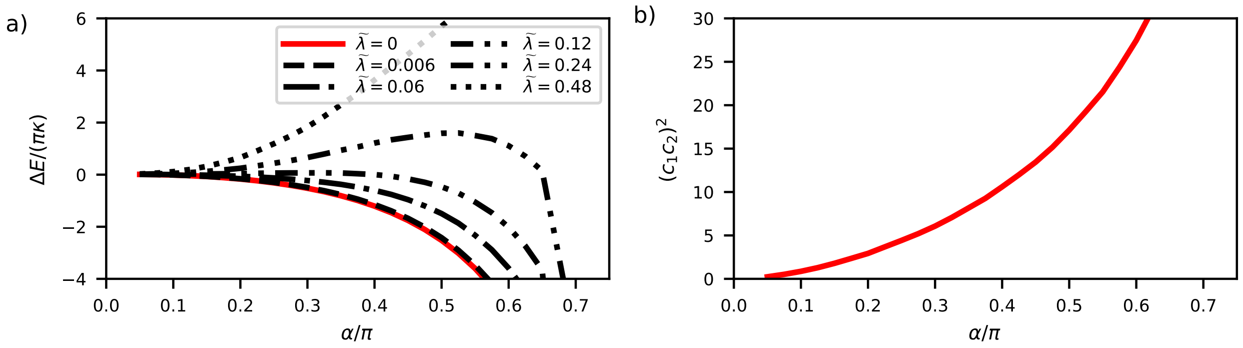


**Figure S6: Line tension** a) Energy (Eq. 6 main text and Eq. S11) for $\epsilon$=2.0 and $\widetilde{\sigma}$=0.1 and different $\widetilde{\lambda}$. b) Squared rescaled Gaussian curvature at the boundary between coated and uncoated region.

### ESCRT-III depletion

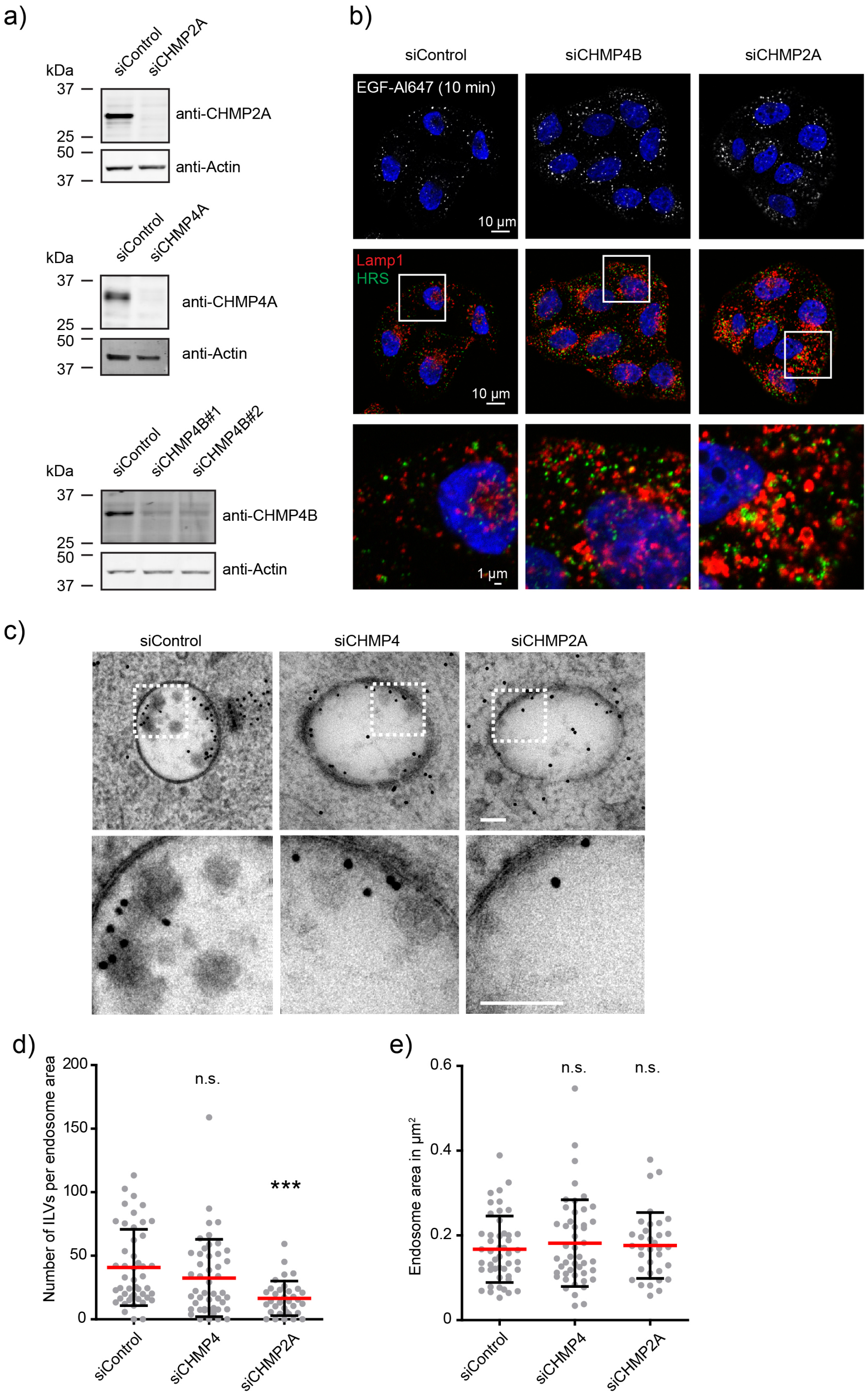


**Figure S7: Characterization of ESCRT-III depleted HeLa cells by immunofluorescence stainings, Western Blot and electron microscopy analysis** a) Western blots showing depletion of CHMP2A, CHMP4A and CHMP4B. CHMP4B siRNA oligo#2 was used in all subsequent experiments. b) Immunofluorescence micrographs showing that ESCRT-III depleted cells can internalize EGF, and that LAMP1 positive endosomes are enlarged and accumulate HRS in ESCRT-III depleted cells compared to control treated cells. c) Electron micrographs showing examples of endosomes from control and siRNA treated cells from 200 nm thick sections. The larger 15 nm gold particles represent fiducial markers and the smaller 10 nm gold particles represent labeled EGFR, which have been internalized following stimulation with EGF for 12 minutes. Scale bar 100 nm. d) Graph representing the quantification of the number of ILVs per endosome area from EM sections as in c. The total number of endosomes are collected from 3 independent experiments (control: n=47 endosomes; siCHMP4: n=47 endosomes) or 2 independent experiments (siCHMP2A: n=31 endosomes). Shown is mean +/- SD. Kruskal-Wallis test with Dunn's Multiple Comparison test. ***: $p<0.001$, n.s., not significant. e) Graph representing the quantification of endosome area from electron micrographs from the dataset in c, d. Shown is mean +/- SD. Kruskal-Wallis test with Dunn's Multiple Comparison test. n.s., not significant. Note that the endosome area is slightly, but not significantly increased in ESCRT-III depleted cells compared to control treated cells. This is in contrast to the enlarged late LAMP1 positive endosomes observed by IF in a), and expected since the gold labelled endosomes investigated by EM represent newly formed EGF-induced endosomes (12 min) that did not yet gain a prominent Class E phenotype.
